# Acute stressors experienced by layer breeders do not affect measures of stress and fear in their offspring

**DOI:** 10.1101/2020.03.23.003301

**Authors:** Mariana R. L. V. Peixoto, Niel A. Karrow, Amy Newman, Jessica Head, Tina M. Widowski

## Abstract

Stressors experienced by layer breeders during egg production can lead to changes in the egg hormone content, potentially impacting their offspring, the commercial layers. Genetic differences might also affect the offspring’s susceptibility to maternal experiences. In this study, we tested if maternal stress affects measures of stress and fear in five strains of layer breeders: commercial brown 1 & 2, commercial white 1 & 2 and a pure line White Leghorn. Each strain was equally separated into two groups: “Maternal Stress” (MS), where hens were subjected to a series of 8 consecutive days of acute psychological stressors, and “Control,” which received routine husbandry. Additional eggs from Control were injected either with corticosterone diluted in a vehicle solution (“CORT”) or just “Vehicle.” Stress- and fear-responses of the offspring were measured in a plasma corticosterone test and a combined human approach and novel object test. Both MS and CORT treatments failed to affect the measured endpoints in the offspring, but significant strain differences were found. The offspring of the white strains showed a higher physiological response compared to brown strains, but the White 2 offspring was consistently the least fearful strain in the behaviour tests. Our study found that the acute psychological stressors experienced by layer breeders did not affect the parameters tested in their offspring and that corticosterone does not seem to be the primary mediator of maternal stress in laying hens. This is highly important, as in poultry production, layer breeders are often subjected to short-term stressors. In addition, we successfully dissociated the physiological and behavioural parameters of stress response in laying hens, showing that increased concentrations of plasma corticosterone in response to stress is not directly associated with high levels of fear.

## Introduction

The organizational structure of commercial layer breeding companies follows a pyramidal model; at the top of the pyramid is a relatively small population of pure line elite stock, followed by the grandparent stock, the parent stock, and the commercial layers at the base (Fig 1). In their lifetimes, layer breeders from the parent stock can produce approximately 115 commercial laying hens (1,2); therefore, a comparatively small number of breeders are needed to provide the total number of commercial layers in the egg system. Parent stocks are typically raised in mixed-sex groups of approximately 100 females per 8 males, either in floor systems housing thousands of birds or in smaller groups in colony breeder cages (2). Similar to the commercial layers, the parent stock may be exposed to stressful procedures and events, such as beak trimming (regulations vary across regions), human handling and social conflicts with other birds.

**Figure 1.**
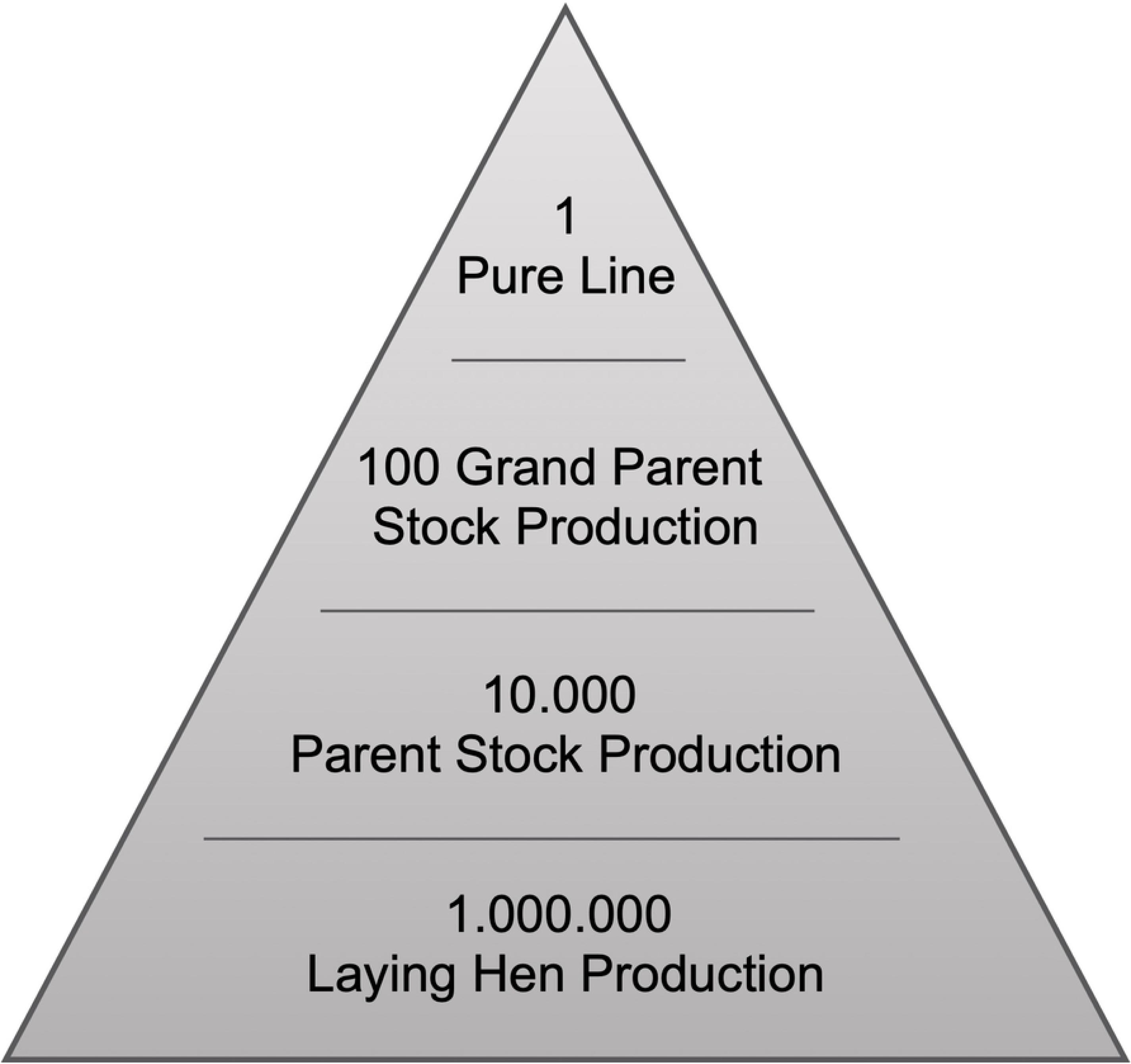
The pyramidal model of layer breeding companies. The organizational structure of commercial layer breeding companies, including the relative number of animals at each level (Modified from (1)).

Once believed that only genetic selection could shape the phenotype of the progeny, it is now known that the over-activation of the hypothalamic-pituitary-adrenal (HPA) axis (i.e., stress response) experienced by the female during egg production can have long-lasting effects in the offspring (3). It has been suggested that these “maternal effects” may shape the phenotype of offspring, which could be adaptive if mother and offspring share similar environments; or detrimental if their environments differ (“environmental matching hypothesis” (3)). Likewise, given that the global population of commercial layers is 7.6 billion birds (4), stressors experienced by layer breeders during egg production may directly affect the growth and survival, and possibly, the overall welfare, of billions of commercial laying hens.

The effects of maternal stress in the offspring of birds and mammals are highly dependent on the intensity, timing of exposure and type of stressor experienced by the mother (5–7). Maternal stress can impact offspring’s physiology, behaviour and cognition (6,8) through structural and functional changes in brain areas involved in the mediation of fear and anxiety (9), as well as through changes in gene expression (10). We have previously shown that chicks from a commercial line of layer breeders that were subjected to daily sessions of psychological stressors during egg production showed a decreased occurrence of anxiety-like behaviour (measured in the total number of distress calls expressed during social isolation) compared to a control group (11). In addition, laying hens subjected to unpredictable feed restriction had chicks that stayed longer in tonic immobility (a measure of fearfulness) and spent less time eating when competing with birds from the control group for access to feed in a novel environment (12). Maternal stress has also been linked to changes in the HPA-axis of the offspring, increasing the concentration of plasma corticosterone in response to capture and restraint in Japanese quails, without changing their baseline hormone concentration (13).

One product of HPA-axis activation during stress in layer breeders is the glucocorticoid hormone corticosterone, which is naturally deposited into the yolk during egg formation (14), and is the exclusive source of glucocorticoids to the embryo until the 16^th^ day of incubation, when the HPA-axis becomes fully functional (15). Due to its pleiotropic effects influencing the expression of thousands of genes, corticosterone has been suggested as a mediator of maternal effects on the offspring (16). Previous studies have shown strong effects on metabolic and developmental processes (6,8,17), but the extent to which corticosterone affects the behaviour of the offspring remains unclear and appears to depend on delivery method and species (18). More recently, other biological components in the egg, such thyroid hormones (19,20), antioxidants (21), immunoglobulins (22), and especially androgens (23) have also been suggested as potential mediators of maternal stress in poultry species.

A laying hen’s susceptibility to maternal stress might also be affected by the different behavioural and physiological responses to stressors observed across strains. In a study conducted with brown and white strains of commercial hybrids, De Haas et al. (24) found that high levels of maternal plasma corticosterone were directly linked to an increase in the occurrence of injurious behaviour (severe feather pecking) in the offspring of Dekalb White but not of ISA Brown hens. Moreover, domestication has increased susceptibility to maternal stress, as seen in a study comparing White Leghorn hens to their ancestor, the Red Junglefowl (25).

Although the literature on maternal stress in poultry species has vastly increased in the last decade (26), this is the first study of its kind to use both pure and commercial hybrid lines of layer breeders to further investigate the interaction between maternal stress, genetics and the behaviour and physiology of the offspring. For this, five strains of layer breeders were equally divided and subjected to either a maternal stress model (“MS”) which involved subjecting the breeder hens to acute psychological stressors; or to a pharmacological stress model (“CORT”), in which corticosterone diluted in a vehicle solution was directly injected into the fertile egg from non-stressed hens (“Control”). A “Vehicle” treatment was included to account for the effects of egg manipulation. Measurements of stress response and fearfulness were assessed in the progeny at 13 and 16 weeks of age, respectively. We predicted that the CORT treatment would show a strong effect in all strains and that the effects of MS would vary according to the natural resiliency of each strain of layer breeder.

## Material and methods

The birds used in this study were treated in accordance with the Canadian Council on Animal Care, and all procedures were approved by the University of Guelph Animal Care Committee (Animal Utilization Protocol #1946). The strains presented herein were anonymized as required by the genetics companies that donated the eggs.

### Parent stock: Management

A total of 2,600 fertilized eggs of 5 strains of parent stock were provided by the University of Guelph’s Arkell Poultry Research Station (pure line White Leghorn) and by two commercial genetics companies: Brown 1 and White 1 from genetics company 1; and Brown 2 and White 2 from genetics company 2. Each company donated 360 female- and 64 male-line hatching eggs per strain.

Eggs from all strains were collected from grandparent hens that were between 40 and 50 weeks of age, subjected to identical incubation and chicks were reared and raised under similar husbandry conditions (17). Chicks were wing-banded at hatch, and each strain was equally distributed into 4 parent flocks that were randomly placed in 2 rooms containing 10 pens of 27 birds (24 females and 3 males) each. Pens (3.7 m^2^) were enriched with pine shavings, one elevated perch and one lower perch, and 5 nest boxes were added at 18 weeks. Flocks were visually separated from each other and did not interact at any moment. Apart from the routine husbandry, all human interaction was avoided to prevent possible habituation.

### Parent stock: Experimental design

Each strain of the parent stock had two pens assigned to the control treatment and two pens assigned to the MS treatment. The control groups were strictly submitted to regular husbandry, while MS hens were subjected to 8 consecutive daily sessions of acute psychological procedures at 3 different ages: 32, 52 and 72 weeks. At the end of the 8^th^ day of stressors, fertile eggs from both treatments were collected, and additional eggs from the control treatment were either injected with corticosterone diluted in vehicle (10ng/mL egg content) (“CORT”) of just vehicle (“Vehicle”). All eggs were incubated, and the offspring flocks from each maternal age were treated as replicates over time (Fig 2). This experimental design allowed us to work with a larger sample size, but it also resulted in replicates confounded with incubatory settings, chick transfer and placement from the incubator to pens, and egg composition, since the nutritional value of the egg changes as a hen ages (27).

**Figure 2.**
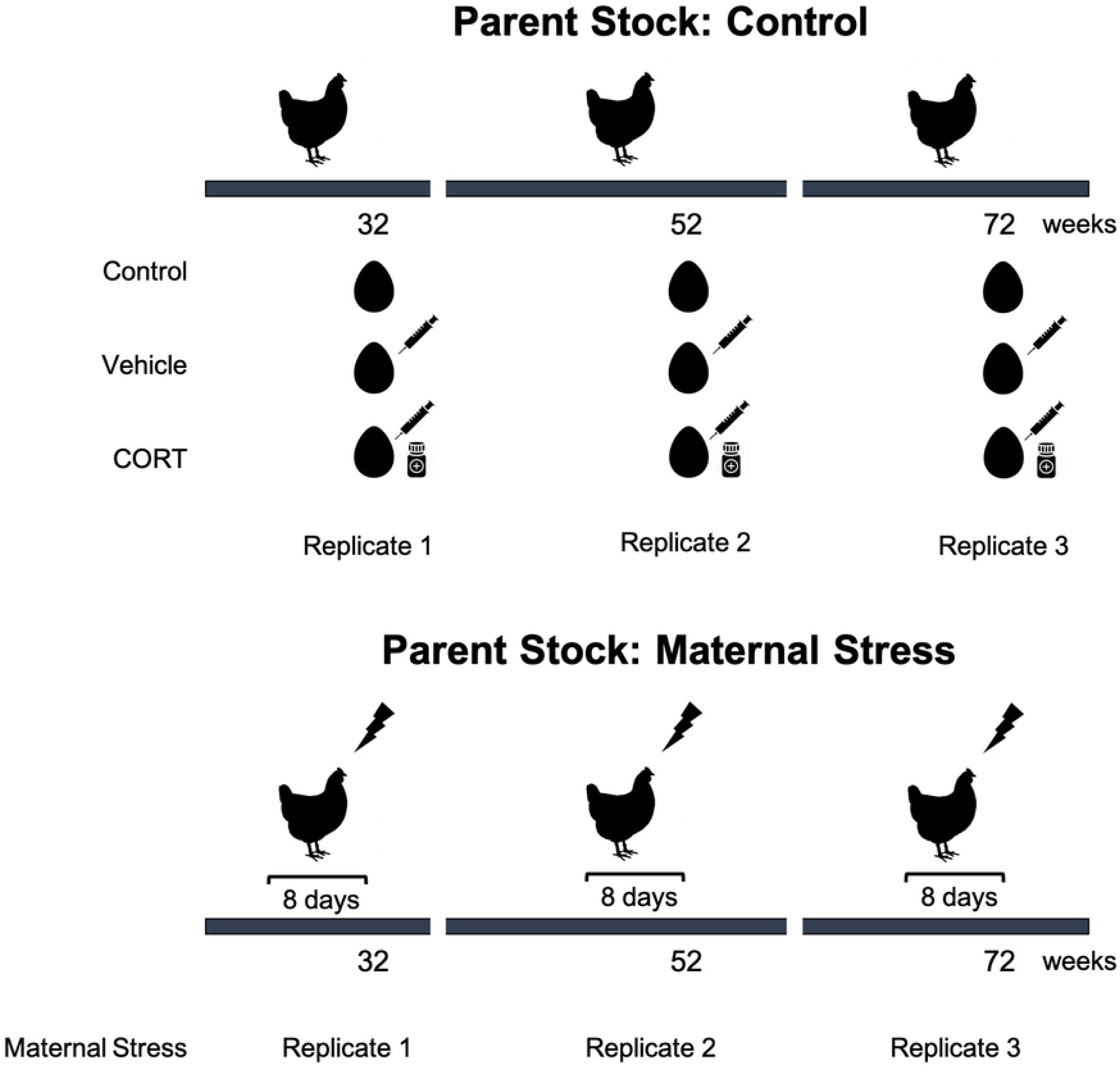
Experimental design. The Control treatment was subjected to regular husbandry, while hens from the Maternal Stress treatment were subjected to 3 sessions of 8 days of stressors prior to egg collection at different ages (32, 52 and 72 weeks). A subsample of eggs from the Control treatment were collected and injected with either corticosterone diluted in vehicle (“CORT”) or just vehicle (“Vehicle”). All eggs were collected and incubated under similar conditions and the different offspring groups were treated as replicates.

### Parent stock: Maternal Stress treatment

The females of the MS flocks were subjected to daily sessions of acute psychological stress procedures that were selected based on their ability to increase plasma corticosterone concentration in avian species (see references for each test below). Since the average time window for egg production from the beginning of vitellogenesis until laying is of approximately 8 days, each MS flock received a minimum of 8 consecutive days of stressors before the beginning of egg collection. Stressors and egg collection were performed until the total number of eggs necessary for incubation had been collected.

Each flock of the MS treatment was subjected twice to the following procedures: 1. Hens were equally distributed into 2 plastic crates (89 cm long × 60 cm wide × 26 cm high) (12 hens / crate), followed by 15 minutes of transportation (28); 2. Hens were individually removed from their home-pens and placed inside a cloth bag located in a nearby room for 10 minutes of physical restraint (29); 3. Hens were crated into 2 groups of 12 birds, transported to an empty room 400 m away from their home-pen and transferred to a test box (100 cm × 100 cm × 200 cm) constructed of solid white panels with 2 doors located on opposite walls and 2 LED lights on the ceiling for 30 minutes. In the box, the hens were exposed to 3 simulations of a predator attack (30 seconds/each) using the silhouette of a sparrow-hawk made of black cardboard (35 cm × 50 cm)(30); 4. Hens were crated and transported to the test box for 15 minutes. During that time, an air horn was blown for 3 seconds every 5 minutes (31,32); 5. Hens were crated, transported to the test room and placed inside the test box for 30 minutes with hens from a different strain (43).

All hens were immediately returned to their home-pens after each stressor. The stress sessions respected the following criteria: 1. All flocks were subjected to one stressor a day; 2. The minimum interval between the application of the same stressor was 4 days to avoid a decrease in the physiological response of the birds due to repeated exposure (i.e., habituation); 3. The stressors were randomly applied from 9:00 to 16:00 h.

### Parent Stock: Vehicle and CORT Treatments

Increased concentrations of maternal plasma corticosterone have been suggested as an efficient tool to increase corticosterone concentration in the egg (14). The CORT treatment aimed to increase the concentration of corticosterone in fertilized eggs from breeder hens. According to previous studies, the basal level of corticosterone in the plasma of laying hens ranges from 0.3 to 5 ng/mL (34), reaching 30 ng/mL in response to stress (35). Previous studies reported that the concentration of corticosterone in egg yolks ranges from 0.77 to 2.8 ng/g in Hy-Line Brown (36– 38) to average 1.6 ng/g in Hy-Line White (36) and 2.13 ng/g in Bovan White (39) under control conditions. The mean concentration of corticosterone in eggs from unstressed birds has been previously reported as 1.17 in yolk and 1.55 ng/mL albumen (40). However, analytical validation of enzyme- and radio-immunoassay techniques showed the presence of cross-reactive substances that hamper quantification of corticosterone in the yolk and albumen of eggs (41); and recent work has shown that even more precise techniques such as Celite or high performance liquid chromatography may not be sufficient to accurately quantify the corticosterone concentration in yolk (reviewed in Groothuis et al., (23)). Therefore, since the exact concentration of corticosterone in eggs from stressed birds remains unknown, we followed the methodology proposed by Janczak et al. (42) and described in Peixoto et al. (17), that was based on plasma corticosterone concentration of stressed hens.

Injections of 10 ng/mL cortisol diluted in sesame oil (“CORT”), or sesame oil alone (“Vehicle”) were used. One day before each egg incubation day, a thin layer of silicone sealant (General Electric, Boston, MA) was applied to a 2 cm × 3 cm area on the basal tip of a sub-sample of control eggs; this sealant prevents gas exchange and contamination following perforation and injection through the shell. On the morning of each incubation day, Vehicle and CORT solutions were prepared. The average weight of egg content, which is estimated to be 90% of total egg weight (43), was 50, 50 and 59 g per hen age group; thus, a volume of 50 μL of either CORT or Vehicle solutions were injected into eggs from 32 and 52 week-old breeders, while 60 μL was injected into eggs from 72 week-old breeders. Injections were performed using a sterile 23-gage needle through a small hole that was perforated through the silicone layer using an egg piercer. Eggs from all treatments were immediately incubated.

### Offspring Stock: Management and data collection

Egg collection, incubation and hatch occurred under similar conditions for all offspring groups. Chicks from each maternal age were individually wing-banded at hatch. The placement of chicks was randomized for each strain and treatment across 40 pens (3.72 m^2^/each) equally distributed in 4 rooms. All pens were enriched with 2 perches and litter floor.

The offspring replicate groups aimed to comprise 2 pens with 20 chickens for each strain and treatment (10 female: 10 male); however, final densities varied due to lower hatchability of the injected eggs (17). The test orders for the procedures described below were balanced across period of the day for all flocks, strains and treatments, in order to minimize the effects of time and circadian rhythm on the results.

### Offspring stock: Plasma corticosterone

To assess the birds’ basal plasma corticosterone level, stress response and stress recovery, 3 consecutive blood collections were drawn from chickens between 13 and 14 weeks of age (N = 675; Table 1). Due to sampling difficulties, fewer samples were collected from birds from the first replicate group, especially the White Leghorn strain.

**Table 1.** Number of birds tested by treatment, strain and replicate for each variable.

| Variable | Treatment | Brown 1 |  |  | Brown 2 |  |  | White 1 |  |  | White 2 |  |  | White Leghorn |  |  |
| --- | --- | --- | --- | --- | --- | --- | --- | --- | --- | --- | --- | --- | --- | --- | --- | --- |
|  |  | Replicate |  |  | Replicate |  |  | Replicate |  |  | Replicate |  |  | Replicate |  |  |
|  |  | 1 | 2 | 3 | 1 | 2 | 3 | 1 | 2 | 3 | 1 | 2 | 3 | 1 | 2 | 3 |
| Stress Response | Control | 7 | 12 | 12 | 12 | 9 | 12 | 12 | 12 | 12 | 12 | 12 | 12 | 8 | 12 | 12 |
|  | M. Stress | 7 | 12 | 12 | 12 | 12 | 12 | 10 | 12 | 12 | 12 | 12 | 12 | 9 | 12 | 12 |
|  | Vehicle | 11 | 12 | 12 | 8 | 12 | 12 | 6 | 12 | 12 | 10 | 12 | 12 | 6 | 9 | 12 |
|  | CORT | 12 | 12 | 11 | 8 | 12 | 11 | 11 | 12 | 12 | 7 | 12 | 12 | 6 | 11 | 10 |
| Total |  | 37 | 48 | 47 | 40 | 45 | 47 | 39 | 48 | 48 | 41 | 48 | 48 | 29 | 44 | 46 |
| Novel Object & Human Approach | Control | - | 4 | 4 | - | 4 | 4 | - | 4 | 4 | - | 4 | 4 | - | 4 | 4 |
|  | M. Stress | - | 4 | 4 | - | 4 | 4 | - | 4 | 4 | - | 4 | 4 | - | 4 | 4 |
|  | Vehicle | - | 4 | 4 | - | 4 | 4 | - | 4 | 4 | - | 4 | 4 | - | 4 | 4 |
|  | CORT | - | 4 | 4 | - | 4 | 4 | - | 4 | 4 | - | 4 | 4 | - | 4 | 4 |
| Total |  | - | 16 | 16 | - | 16 | 16 | - | 16 | 16 | - | 16 | 16 | - | 16 | 16 |

Following the methodology proposed by Wingfield (44) and replicated by Goerlich et al., (45), selected birds were individually captured from their home pen, and a baseline blood sample was drawn from the wing vein within 3 minutes from capture. Following, the bird was placed inside a large cloth bag for 10 minutes and the second blood sample (“stress-induced response”) was collected. After 20 more minutes of physical restraint, the last sample was collected. All samples were centrifuged for 15 minutes with equipment settings at 2,500 RPM, 4.2 rotor at 4°C. Duplicates of each plasma sample were collected and stored at −4°C until analyses. Total glucocorticoid concentration in plasma was determined using a commercial corticosterone enzyme-linked immunosorbent assay (ELISA) kit (Enzo Life Sciences, NY, USA). Samples were tested in duplicate following a standard protocol (see the online manual: http://www.Enzolifesciences.com/ADI-900-097/corticosterone-eia-kit/). Our assay had a sensitivity of 21.75 pg/ml and 13.6% and 6.0% inter-assay and intra-assay coefficients of variation, respectively.

### Offspring stock: Human approach and novel object test

Fear behaviour can be quantified by measuring the conflicting motivation to approach or avoid a potentially dangerous stimulus (46). In the human approach (HA) and novel object (NO) test, we measured the duration of time that the birds spent in zones located a distance away from either a person and a novel object (umbrella) (47). The longer the duration of time spent away from the stressor, the more fearful the bird was considered to be.

Birds at 16 weeks of age (N = 160; Table 1) were tested in same-sex pairs. Testing was conducted in an arena located in a quiet room close to the birds’ home-pens. The arena measured 210 cm × 180 cm × 120 cm and contained a 50 cm × 50 cm × 70 cm habituation box at one end and a clear netting wall at the opposite side, from which the animals could see the stressor (person or NO) positioned 20 cm outside it. The floor was separated into two equal parts (Zone 1 and Zone 2) that were used to determine the birds’ location (Figures 3 and 4).

**Figure 3.**
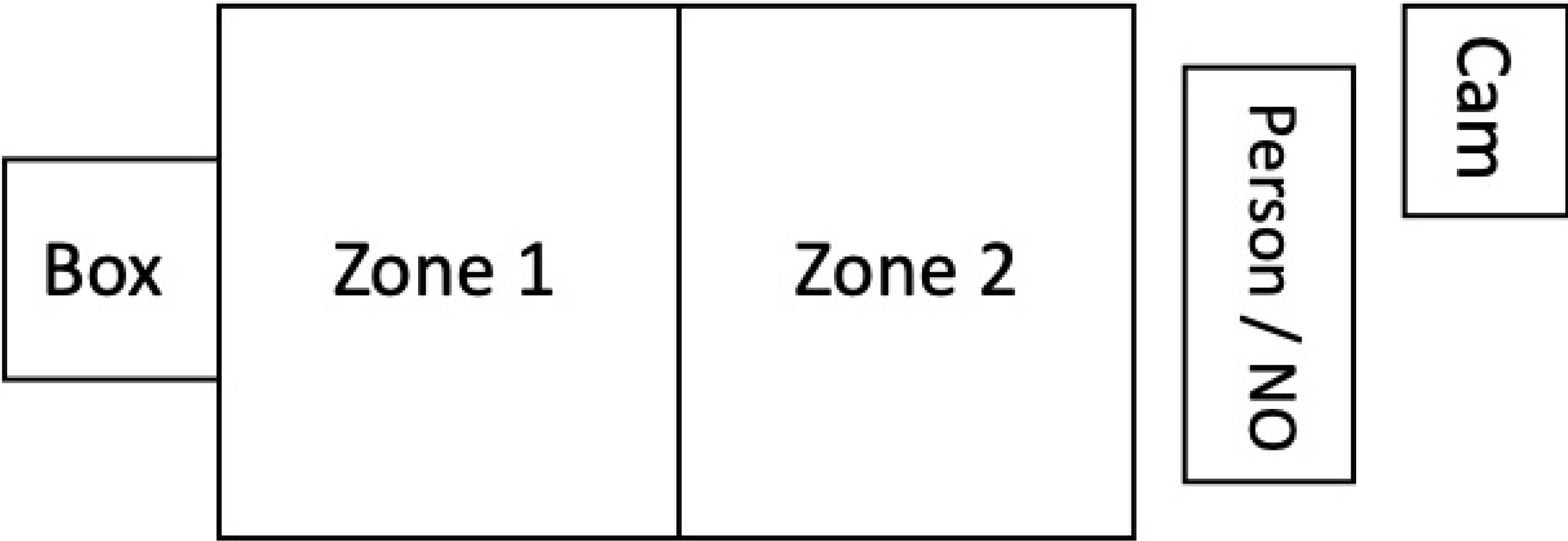
Illustration of the test arena used in the human approach and novel object tests. Following a modified version of Brantsæter et al. (47), birds were placed in the habituation box (“Box”) with access to the arena (“Zone 1” and “Zone 2”) for 30 seconds. Then, with a person positioned at the other end of Zone 2, birds were given access to Zone 1. After 5 minutes, the person was replaced by an umbrella (NO). Recordings were done using a security camera, placed above the middle of the arena in the ceiling and a camcorder (Cam), located adjacently to the stressor.

**Figure 4.**
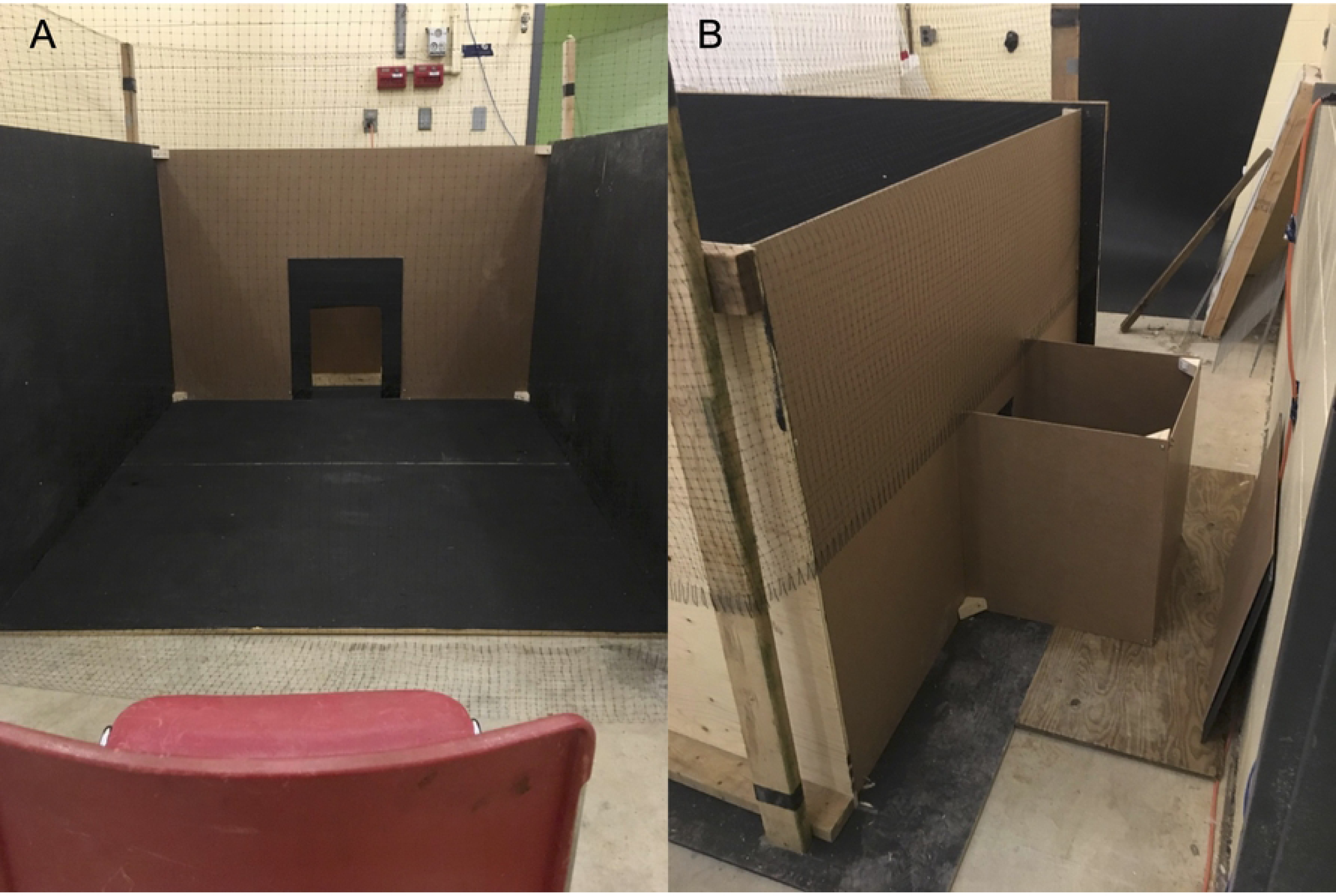
Human approach and novel object arena. A. Front view of the arena. B. Close-up of the habituation box positioned behind Zone 1.

The birds were marked with livestock paint two days before testing to allow for individual identification on video analyses. On the test day, each pair of animals was quietly removed from their home-pen and moved into the start box attached to the arena. After 30 seconds, a pull-up door was opened, giving the birds access to the arena. At this moment, the “observer” (a person who had not been seen by the birds on that day), was sitting on the chair, facing the arena, but avoiding direct eye contact with the animals. We used a total of three different observers, all women wearing a dark blue coverall. After 5 minutes in front of the animals, the observer opened a pink umbrella pointing to the arena, placed the object onto the chair, and quietly left the room, thus starting the novel object test, which also lasted 5 minutes. To record both tests, we used one camcorder (Panasonic HC-V180K) positioned adjacently to the stressor and one security camera (Panasonic WV-SPN531) attached to the ceiling, facing the centre of the arena.

Measurements of behaviour were analyzed from the video recordings by one observer blinded to treatment and strain. Chickens within pairs were individually identified based on their paint marking and their behaviour was quantified. Latency to leave the start box (s) and time spent in each zone (s) was assessed through continuous observation of both HA and NO tests. A bird’s location was determined if both of the animal’s feet were in the same zone. Since bird location varied at the beginning of the NO test,, we recorded where the birds were positioned when the umbrella was opened; and for those located outside of the box, their flight response to the umbrella (flight, freeze, walk; Table 2) (N = 132). Due to external factors, data from replicate 1 were not included in the analyses. All birds were immediately returned back to their home-pens after testing.

**Table 2.** Ethogram used in the human approach and novel object behaviour measurements.

| Behaviour | Description |
| --- | --- |
| Fly | Bird jumps or initiate flying directed to surrounding, avoiding the stimulus |
| Freeze | Bird stands upright and immobile for a minimum of 5 seconds |
| Walk | Two or more steps, attending to surroundings, head erect |

### Statistical analyses

The Glimmix procedure of SAS 9.4 (SAS Institute, Cary, NC) was used to perform all statistical analyses. The basic statistical model included fixed effects of treatment, strain, sex, and a treatment by strain interaction. Random effects were grouped by strain and included offspring group and pen nested in room. Further pre-planned comparisons included treatment (Control versus Maternal Stress and Control versus CORT), strain (white versus brown lines) and Genetic Company 1 (Brown 1 and White 1) versus Genetic Company 2 (Brown 2 and White 2). Tests for normality included Shapiro-Wilk and Anderson Darling measurements in conjunction with visual plots. When a positive interaction was found, analyses controlled for the multiple testing error using the percentage of false-positives, which estimates the false discovery rate (48). Significance was declared at P < 0.05.

### Plasma corticosterone

The basic model in ANOVA was adjusted for repeated measures, and the following fixed effects were added: timepoint (baseline, 10 minutes and 30 minutes), timepoint by strain, and timepoint by treatment. The 3-way interaction “strain by treatment by timepoint” was not statistically significant (P = 0.755) and was therefore removed from the model. Random effects were grouped by strain and included offspring group and pen nested in room, with bird as the experimental unit. Data were assumed to be distributed according to a log-normal transformation. Significance post-FDR correction was set at P < 0.01. Least Square (LS-) means and standard error of means (SEM) were back-transformed and are presented in the results as the average concentration of corticosterone in plasma (ng/ml).

### Human approach and novel object

Time spent in the arena was analyzed in ANOVA and results were partitioned by test. Fixed effects included the basic model plus zone (Box, Zone 1 and Zone 2), strain by zone, treatment by zone and sex by zone. The 3-way interaction “strain by treatment by zone” was not statistically significant (NO: P = 0.078; HA: P = 0.067) and was therefore removed from the model. Random effects were grouped by strain and included offspring group, pen nested in room and location in the arena at the beginning of each test; individual bird (nested in pair) was the experimental unit. To meet the assumption of a normal distribution, data were log-normally transformed. Significance post-FDR correction was set at P < 0.02. LS-means and SEM were back-transformed and are presented in the results as the average time (s) spent in each zone. The percentage of birds that spontaneously entered the arena and the type of response to stressor at the moment when the umbrella was opened were analyzed using a Chi-square test. Results are presented as the percentage of each response by strain.

## Results

### Plasma corticosterone

To measure if strain, treatment or sex affected corticosterone concentration in blood plasma at different timepoints, we specifically aimed for interactions between these effects and the time of collection (“timepoint”). The offspring from different treatments showed no differences in their plasma corticosterone concentration at any time of collection (P = 0.835), but the average of corticosterone produced throughout the test varied among strains (P = 0.010) (Table 3). As expected, corticosterone concentration increased in all strains 10 minutes after the onset of testing (P < 0.001), but only the white strains showed a decrease in hormone concentration after 30 minutes of restraint (Figure 5).

**Table 3.** Physical restraint test. Effects of sex, strain, treatment and timepoint on plasma corticosterone concentrations and contrast comparisons by treatment and strain.

| Stress Response |  |  |
| --- | --- | --- |
| Effect |  | P-value |
| Sex |  | 0.156 |
| Strain |  | < 0.001 |
| Treatment |  | 0.102 |
| Timepoint |  | < 0.001 |
| Strain x Treatment |  | 0.202 |
| Strain x Timepoint |  | 0.010 |
| Treatment x Timepoint |  | 0.835 |
| Contrasts |  |  |
| Treatment | Control vs Maternal Stress | 0.745 |
|  | Control vs Vehicle | 0.717 |
|  | Control vs CORT | 0.693 |
| Strain | White vs Brown | < 0.001 |
|  | Genetic Company 1 vs 2 | 0.596 |

**Figure 5.**
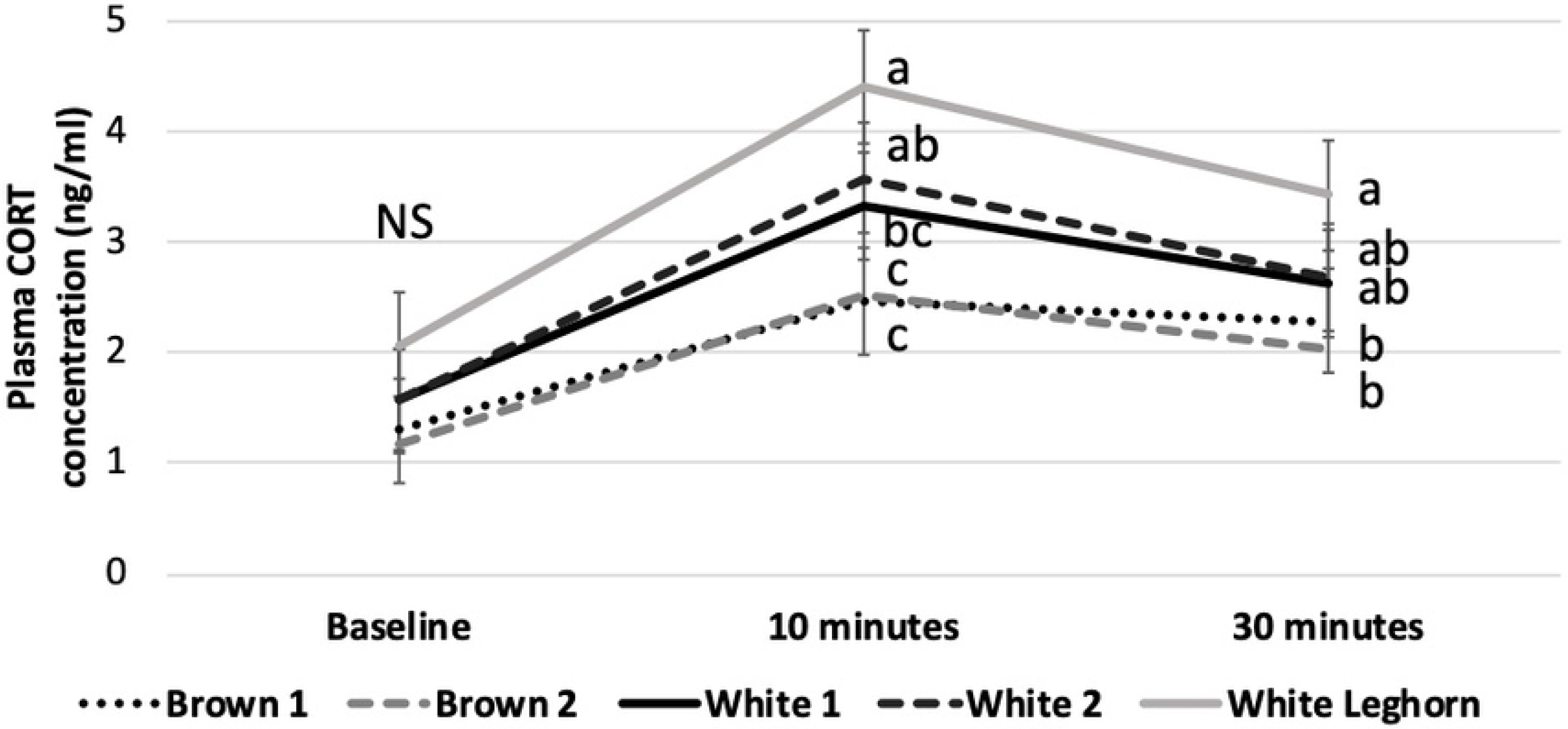
Plasma corticosterone. Mean corticosterone (± SEM) at baseline, 10 and 30 minutes after onset of physical restraint, displayed by strain. Means with different superscripts (a-c) within the same timepoint differ (P < 0.01).

The baseline corticosterone concentration was similar for all strains; however, the White Leghorn offspring consistently showed the highest concentration of circulating corticosterone after 10 and 30 minutes of physical restraint, while both commercial brown strains displayed the lowest values at the same timepoints (Figure 5). Post-hoc contrast analyses indicate a difference between the hormone concentration of brown- and white-coloured strains (P < 0.001).

### Human approach and novel object test

Similar to plasma corticosterone, we specifically tested for interactions between a bird’s location in the arena (“Zone”) with strain, treatment and sex in order to find if time spent in different locations was affected by these variables. In the human approach test, the duration spent at the arena zones was not affected by treatment or sex but varied according to genetic strain (Table 4). The Brown 2 offspring displayed the highest degrees of fear in HA, spending more time in the start box and less time close to the observer than the other strains. In contrast, the White Leghorn offspring showed the shortest latency to leave the box, while the White 2 offspring spent the longest duration closer to the person (“Zone 2”). Once in the arena, the birds did not return to the habituation box in any of the tests.

**Table 4.**
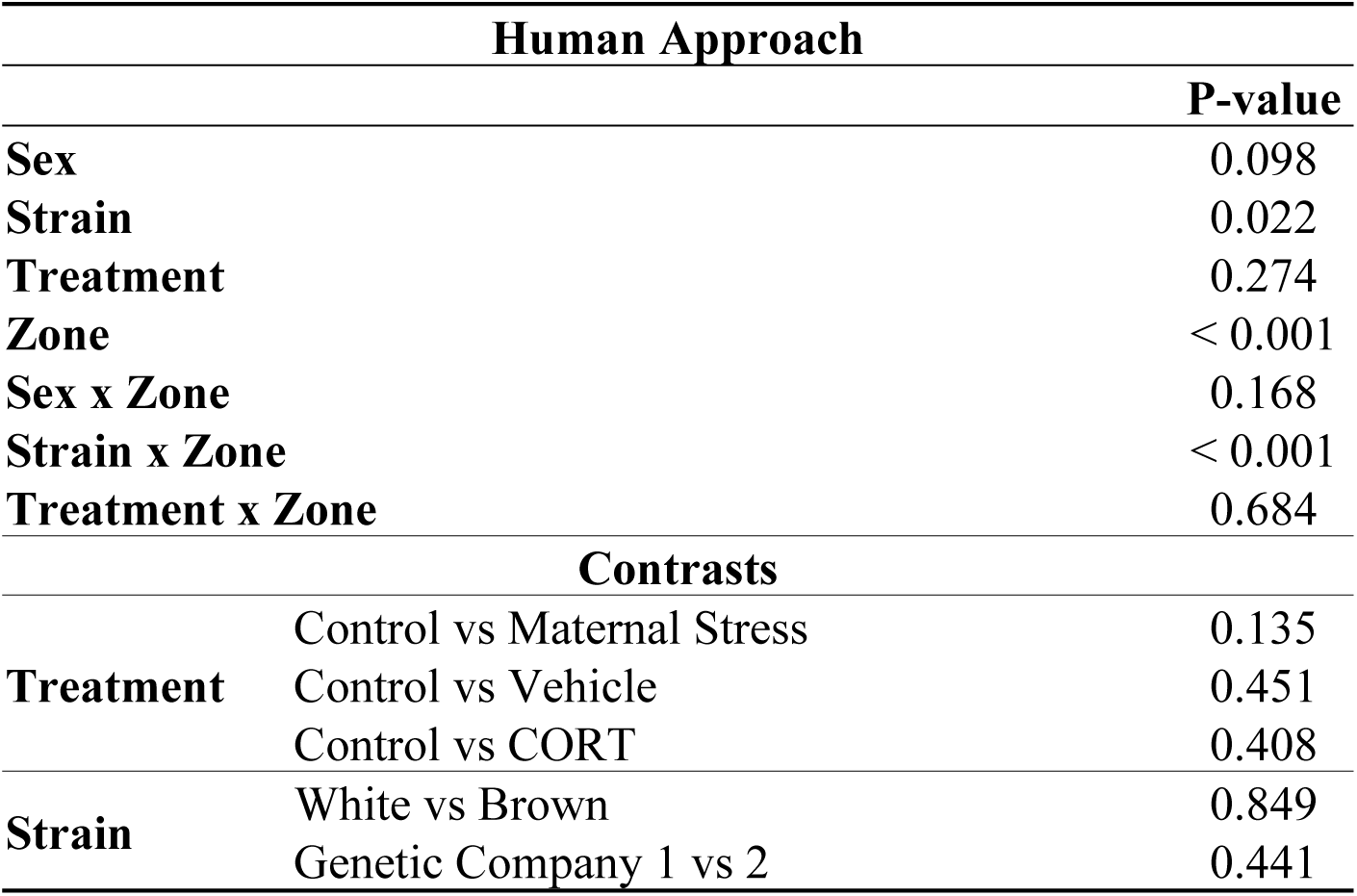
Human approach test. Effects of sex, strain, treatment and arena zone (“Zone”) on time spent in the arena and contrast comparisons by treatment and strain.

Treatment failed to affect the duration of time spent in different arena locations (zones) during the novel object test; however, strain differences were found (Table 5). All strains spent similar durations inside the box and in zone 1, but the White Leghorn offspring spent the shortest duration in zone 2, standing out as the most fearful strain in the presence of a novel object. Females showed a higher latency to leave the habituation box and spent less time close to the novel object than males (Figure 8).

**Table 5.**
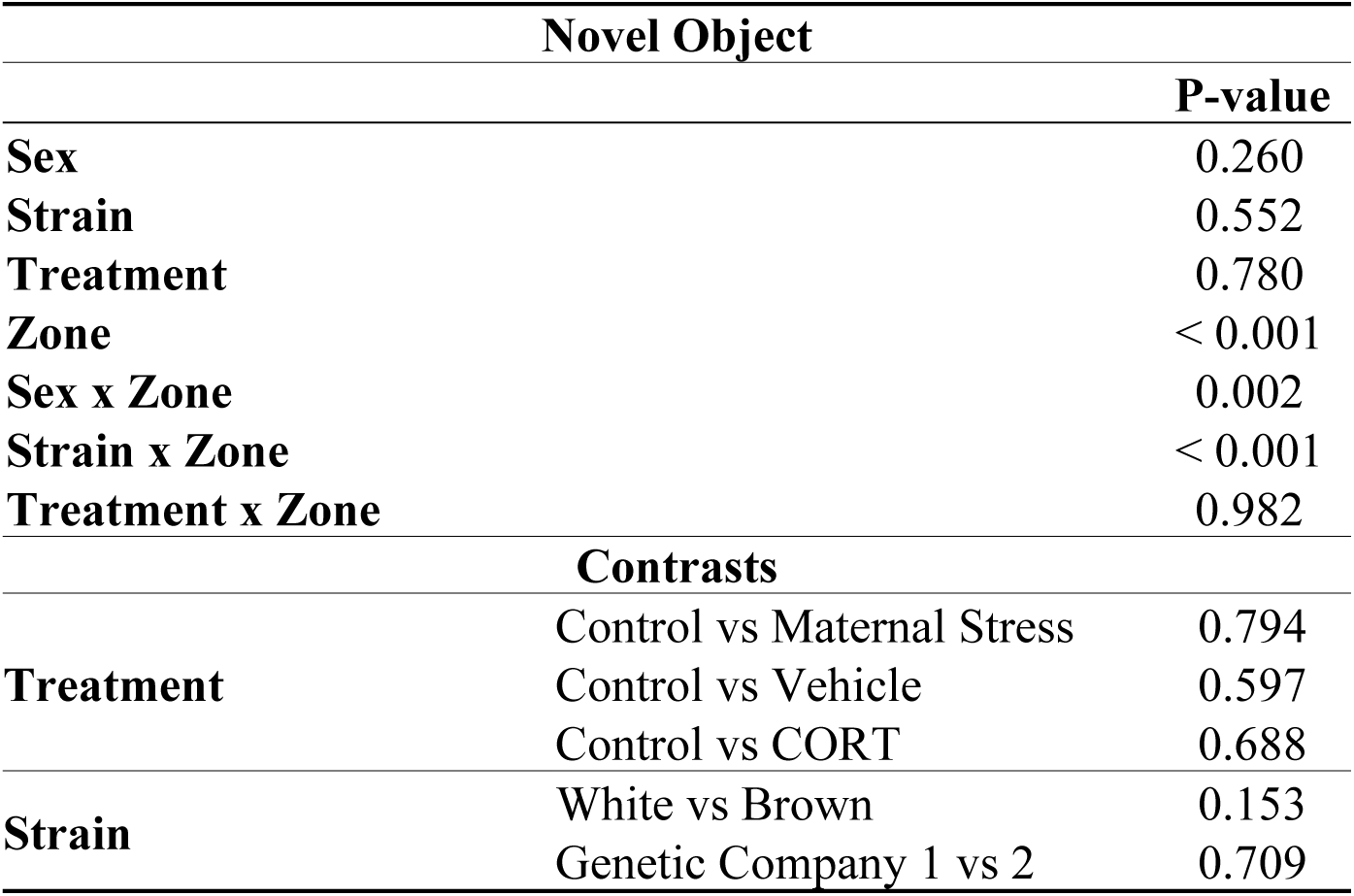
Novel object test. Effects of sex, strain, treatment and arena zone (“Zone”) on time spent in the arena and contrast comparisons by treatment and strain.

**Figure 6.**
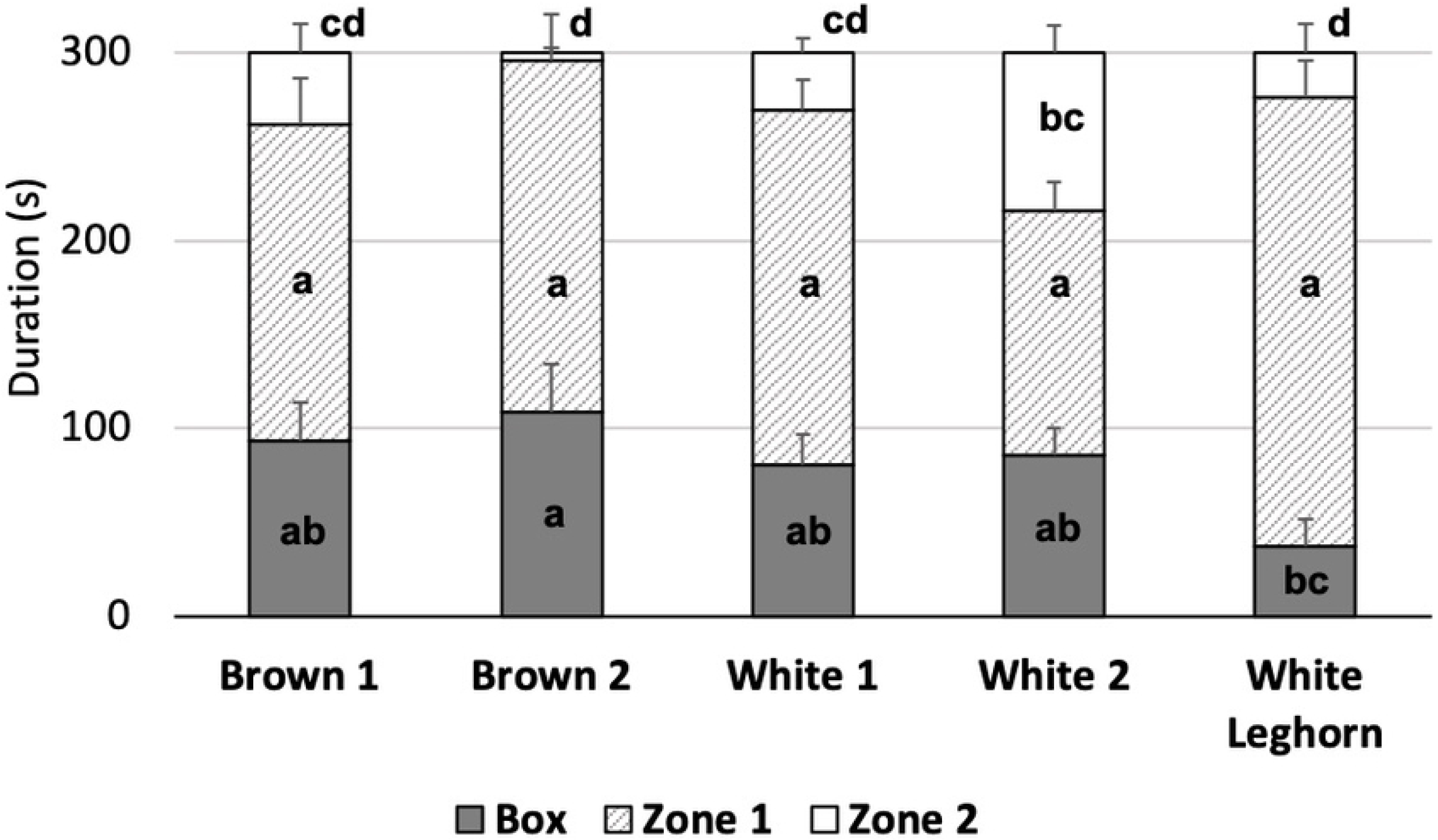
Human approach test. Average time (s) spent in different arena locations (zones) displayed by strain (± SEM). Statistical differences in duration of time spent in different zones across strains are indicated by letters (a-d) (P < 0.02).

**Figure 7.**
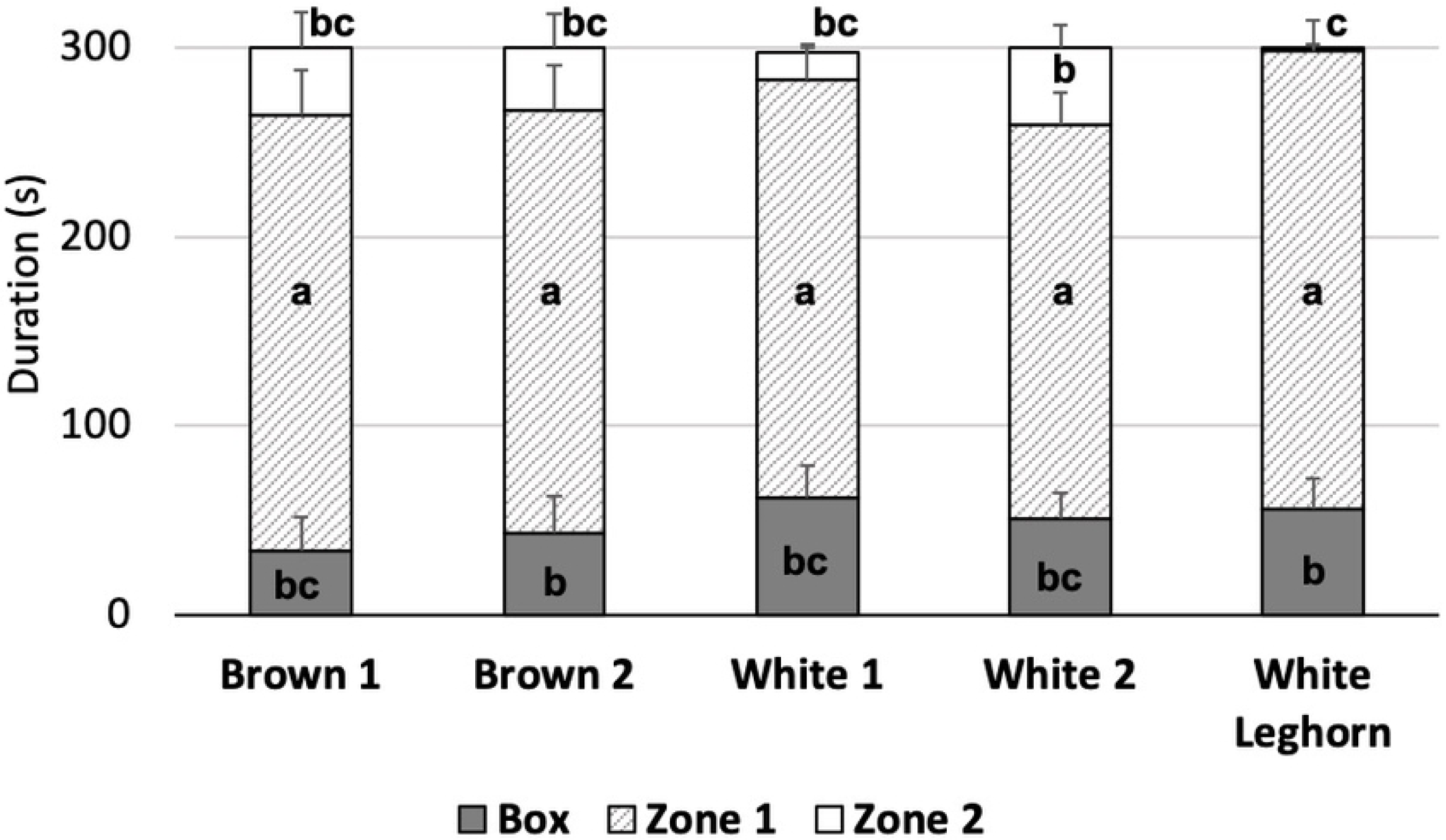
Novel object test. Average time (s) spent in different arena locations displayed by strain (± SEM). Statistical differences in duration of time spent in different zones across strains are indicated by letters (a-c) (P < 0.02).

**Figure 8.**
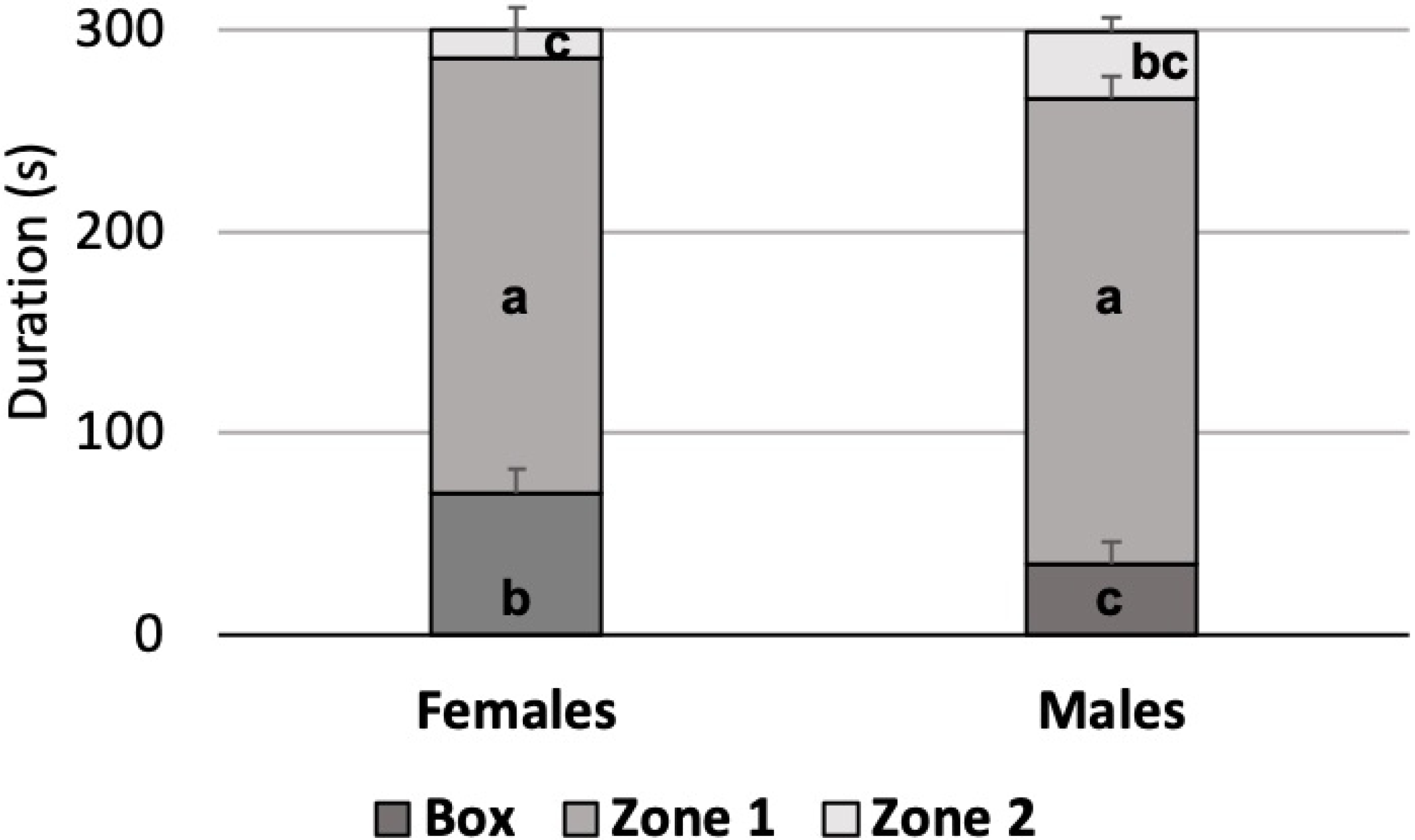
Novel object test. Average time (s) spent in different arena locations displayed by sex (± SEM). Statistical differences in duration of time spent in different zones between females and males are indicated by letters (a-c) (P < 0.02).

Descriptive statistics of the birds’ location at the moment when the umbrella was opened are presented in Table 6, with 61.9% in Zone 1. Behaviour response to sudden movement (umbrella opening) varied among strains; commercial- and pure line-white strains flew away from the umbrella more frequently than both brown strains (Fig 9).

**Table 6.** Location of the birds at the moment when the umbrella was opened.

| Strain | Location |  |  |  |  |  |
| --- | --- | --- | --- | --- | --- | --- |
|  | Box |  | Zone 1 |  | Zone 2 |  |
|  | # | % | # | % | # | % |
| Brown 1 | 7 | 22.5 | 18 | 57.0 | 7 | 22.5 |
| Brown 2 | 6 | 17.0 | 22 | 70.0 | 4 | 13.0 |
| White 1 | 5 | 15.5 | 20 | 62.5 | 7 | 22.0 |
| White 2 | 4 | 14.0 | 15 | 47.0 | 13 | 39.0 |
| White Leghorn | 7 | 22.0 | 24 | 76.0 | 1 | 2.0 |
| Total | 29 | 18.1 | 99 | 61.9 | 32 | 20.0 |

**Figure 9.**
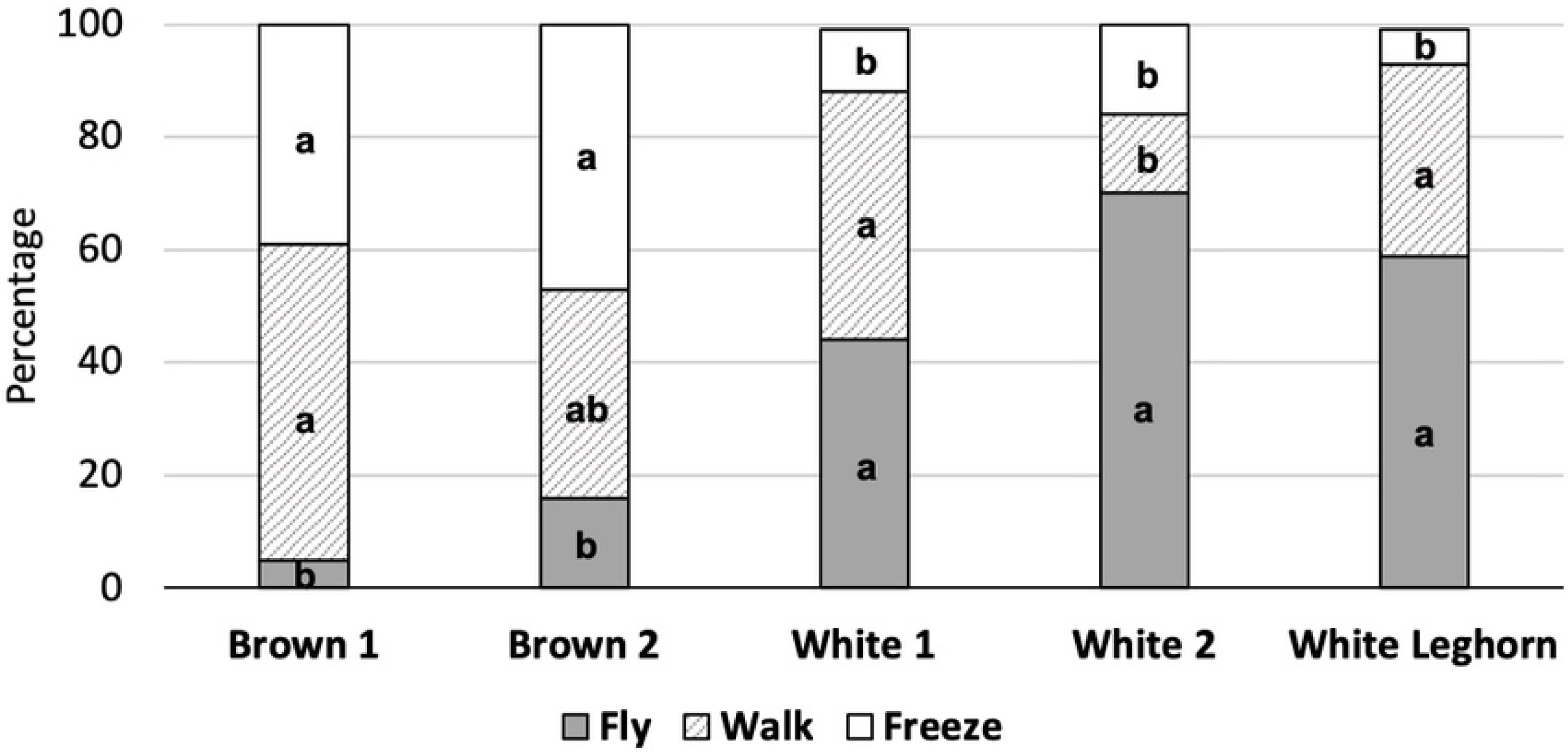
Behaviour response to opening umbrella. Percentages are displayed by strain. Statistical differences within strain are indicated by letters (a,b) (P < 0.05).

## Discussion

### Treatment, strain and sex effects

In this study, we examined the effects of maternal stress on the stress response and fearfulness of the offspring from different strains of layer breeders. We predicted that CORT would show a strong effect in all strains and that the effects of MS would vary according to the natural susceptibility of each strain. None of the stress treatments affected the traits measured, but strain and sex differences were observed in the progeny.

Our results show that the acute stressors experienced by layer breeders during egg production did not shape the behaviour or the stress response of commercial layers. Previous work in poultry that subjected females to long-term, chronic stressors, report carry-over effects on behaviour and physiology of offspring (13,31,49), however ours is the first test of acute stress. Moreover, the mothers of the birds tested in the previous studies were housed in cages, a contributing factor to the occurrence of maternal effects (50). Our work is highly important, as in poultry production, breeders are often subjected to acute stressors (such as handling, vaccination and loud noises) and these data suggest that these experiences do not seem to affect the offspring.

The lack of results in the CORT treatment suggests a weak or inexistent biological link between maternal corticosterone and the offspring’s phenotype, as previously reported in a passerine species (18). It is possible that the corticosterone concentration used in this study was not sufficient to shape the phenotype of the offspring. However, it is most likely that puncturing and injecting the fertile egg moments prior to incubation, as well as applying a silicon sealant onto the egg surface, are highly invasive and unnatural procedures, which might have exposed the progeny to an additional stressor and unintentionally selected a subset of birds that were more resilient to the adverse effects of the injection; thus, limiting the generalization of these results.

Although no treatment effects were observed, our study revealed significant differences in both behavioural and physiological measures among strains. Results show that the white strains produced more corticosterone in response to physical restraint, approached the person and the novel object more frequently and showed greater flight response (avoidance behaviour) when startled by the umbrella than the brown strains.

Behavioural and physiological responses to stress are commonly interpreted along a proactive-reactive continuum (51). Proactive animals tend to produce lower physiological response to stress (e.g., corticosterone) and have a fast response to a novel stimulus (e.g. fast approach, more aggressive); whereas reactive animals produce higher physiological response to a stressor and display a slow and shy behaviour response (e.g., slower approach, more passive towards a stimulus) (52,53). Our study showed that all white strains produced more corticosterone in response to physical restraint, suggesting a reactive physiological profile. However, the White 2 strain approached both novel stimuli more rapidly than the other strains and displayed a faster avoidance response to the startling stimulus, thus characterizing a proactive behavioural profile.

Mammals and passerines typically display the same profile for both behavioural and physiological responses to stress, thus characterizing the concept of “coping style” (51). In agreement with previous studies (36,53–56), our results show that this concept does not apply to laying hens. Although the majority of birds from the White 2 strain flew away from the umbrella and showed a high concentration of corticosterone in response to physical restraint, they consistently spent more time close the human and the novel object, standing out as the least fearful strain in the behaviour tests. In a previous publication, we also reported that these same white strains produced fewer distress calls in response to social isolation and spent less time in tonic immobility, a measure of fearfulness in chickens, compared to the brown strains (11). Our results, thus, suggest a dissociation between stress reactivity and fearfulness in laying hens.

In many species, physiological traits are associated with pigmentation (57,58) and melanism has been shown to signal the ability to cope with elevated stress hormones in barn owls (59). Effects on aggressive behaviour have also been found (60), but to our knowledge, no studies have yet focused on feather pigmentation and fear-response in birds. Nevertheless, another interesting finding is that the offspring of the Brown 2 strain showed higher degrees of fear of humans, while the offspring of the White 1 and White Leghorn strains displayed a higher response in the presence of the novel object, suggesting that white strains of laying hens might be more fearful of objects than the brown strains. It is important to mention that the greater distances from the novel object observed in the white strains are most likely related to these strains’ response to the umbrella opening. Nonetheless, previous studies have shown that White Leghorn hens were found to be more fearful of a novel object than hens from a Rhode Island Red strain (61), and a positive correlation between fear of a novel object and low body weight in chickens was found (white strains are commonly lighter than browns) (56). Inconsistent results are found in the measurement of fear of humans, possibly depending on other environmental factors, such as husbandry, level of exposure to humans and previous experiences (56). Our findings, thus, suggest that the occurrence of fear-like behaviour in a laying hen depends on the interaction between her genetics and the source of fear.

Lastly, we observed that cockerels spent more time close to the umbrella in the novel object test, displaying lower degrees of fear than the hens in that test. The behaviour of animals are mediated by two gonadal hormones, androgen and estrogen (62). Individually and combined, these hormones act on the neural system, organizing the neuronal circuitry involved in behavioural functions (62,63), thus explaining the occurrence of sex differences in behavioural traits. Sexual dimorphism on fear-related behaviour can also be linked to genetic selection for productive traits through pleiotropic effects; a quantitative trait loci (QTL) related to fear response in the novel object test was found to share positions with a major QTL for growth and body weight in cockerels, but not in hens (64).

### Limitations, challenges and future opportunities

The physical restraint procedure is a well validated test (39) that was used twice in this study: as a stressor in the parent stock and as a measure of stress response in the offspring. As previously reported (11), the layers from the MS treatment were still physiologically responsive to the stressor even after repeated exposure, therefore, validating the physical restraint test as a stressor in the MS treatment. However, no further stressors applied onto the breeder flock in the MS treatment were validated, thus limiting the discussion on whether the lack of effects in the progeny of MS breeders is due to a natural resilience of the breeder hens to the chosen stressors, or to an embryonic resilience to increased hormone levels deposited into the egg by the stressed breeder. Moreover, only the total concentration of corticosterone was measured in the study; but glucocorticoids can be found in both biologically active and inactive forms in the blood. Biologically active (i.e., free) corticosterone is able to bind to receptors within the tissues, whereas glucocorticoids bound to a protein carrier (predominantly corticosteroid-binding globulin (65)) are inactive. To accurately quantify the potential biological activity of corticosterone in plasma samples, it would be necessary to know the concentration of free corticosterone in comparison with total corticosterone (reviewed in 45).

A challenge faced in this study is that the actual concentration of corticosterone transferred from the mother to egg remains largely unknown (Rettenbacher et al., 2009, 2013a; Almasi et al., 2012) and may differ across strains (Navarra and Pinson, 2010). Although analytical techniques such as Celite and HPLC are more precise than the often used enzyme- and radio-immunoassay, they may still not be sufficiently accurate to quantify the concentration of corticosterone in the yolk (reviewed in Groothuis et al., (23)). Therefore, the hormone concentration used in our injections might have been lower than the physiological range of different strains of breeder hens, thus explaining the negative findings. In follow-up studies it would be important to explore any strain-related differences in corticosterone levels transfer to eggs. Moreover, we strongly encourage further investigation other biological mechanisms different than corticosterone. the Currently, the use of hormone injections in fertile eggs as a model for maternal stress remains questionable.

## Conclusion

This research successfully shows that acute stressors such as brief transportation, physical restraint or loud auditory noise experienced by layer breeders during egg production, do not shape the behaviour or the stress response of their offspring, the commercial layers. This finding has significant importance in the agricultural field, as layers breeders are constantly subjected to different types of acute stressors in their lifetimes.

Our results also suggest the disconnection between stress reactivity and fearfulness in laying hens, in which high plasma concentration of corticosterone should not be associated with a more fearful personality. In addition, the occurrence of fear-like behaviour in layers depends on their genetics and the source of fear, varying in response to the source of stimuli. The implications of this study in both scientific and agricultural fields are numerous, as variations in behavioural and physiological responses to stress can be decisive when assessing different strains of layers, or when determining the overall adaptability of a strain to a specific housing system; therefore, having significant welfare and economic impacts in the egg production system.

